# Evaluating the Simple Arrhenius Equation for the Temperature Dependence of Complex Developmental Processes

**DOI:** 10.1101/2020.07.17.208777

**Authors:** Joseph Crapse, Nishant Pappireddi, Meera Gupta, Stanislav Y. Shvartsman, Eric Wieschaus, Martin Wühr

## Abstract

The famous Arrhenius equation is well motivated to describe the temperature dependence of chemical reactions but has also been used for complicated biological processes. Here, we evaluate how well the simple Arrhenius equation predicts complex multistep biological processes, using frog and fruit fly embryogenesis as two canonical models. We find the Arrhenius equation provides a good approximation for the temperature dependence of embryogenesis, even though individual developmental stages scale differently with temperature. At low and high temperatures, however, we observed significant departures from idealized Arrhenius Law behavior. When we model multistep reactions of idealized chemical networks we are unable to generate comparable deviations from linearity. In contrast, we find the single enzyme GAPDH shows non-linearity in the Arrhenius plot similar to our observations of embryonic development. Thus, we find that complex embryonic development can be well approximated by the simple Arrhenius Law and propose that the observed departure from this law results primarily from non-idealized individual steps rather than the complexity of the system.

## Introduction

For more than a century the Arrhenius equation serves as a powerful and simple tool to predict the temperature dependence of chemical reaction rates (Arrhenius, 1889). This equation, named after physical chemist Svante Arrhenius, posits that the reaction rate (k) is the product of a pre-exponential factor A and an exponential term that depends on the activation energy (Ea), the gas constant (R), and the absolute temperature (T).

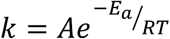

In the late 19^th^ century scientists proposed many relationships between reaction rates and temperature (Berthelot, 1862; Harcourt and Esson, 1895; van’t Hoff, 1893; Schwab, 1883; Van’t Hoff, 1884). Among them the Arrhenius equation stood out at least partly due to how it can be intuitively interpreted based on the later developed transition-state theory (Evans and Polanyi, 1935; Eyring, 1935; Keith J. Laidler and M. Christine King, 1983). Based on this theory, the exponential term of the Arrhenius equations is proportional to the fraction of molecules with greater energy than the activation energy (Ea) needed to overcome the energetically unfavorable transition state for a reaction. The pre-exponential “frequency” factor A is proportional to the number of molecular collisions with favorable orientations.

More recently, the Arrhenius equation’s use has also been extended to more complex biological systems such as the cell cycle duration (Begasse et al., 2015; Falahati et al., 2020), or by extension the Q_10_ rule in proliferation dynamics in populations of bacteria (Martinez et al., 2013). This broad applicability of the Arrhenius equation to complex biological systems is surprising given that these systems involve a myriad of reactions, presumably each with its own activation energy and temperature dependence.

One of the most complicated biological processes the Arrhenius equation has been applied to the development of a single fertilized egg into the canonical body plane of an embryo (Chong et al.). Embryos of most species develop outside the mother and many have evolved so that they can adopt to fairly wide temperature ranges. Canonically, it has been observed that embryos develop faster with higher temperature (Khokha et al., 2002; Kuntz and Eisen, 2014; Sin et al., 2019). However, some proteins decrease their concentration and presumably their activity in the liquid-phase separated nucleolus with increasing temperature (Falahati and Wieschaus, 2017). This is expected as most intramolecular interactions weaken with higher temperature (Ball and Key, 2014). Recently, it has been proposed that the developmental progression of fly embryo development scales uniformly with temperature (Kuntz and Eisen, 2014), which would be only consistent with the Arrhenius equation if all activation energies for rate-limiting transition states are similar. In this case, coupled chemical reactions would collapse into a common Arrhenius equation with one master activation energy and integrated frequency factor. Is it possible that evolution has led to such uniform activation energies in embryos to enable canonical development over a broad temperature range?

To investigate these remaining puzzles and apparent discrepancies, we observed the temperature dependence of developmental progression of fly and frog development. We find that apparent activation energies of different developmental stages vary significantly i.e. the time it takes for embryos to develop through different stages scale differently with temperature. Nevertheless, we show that the Arrhenius equation still provides a good approximation for the temperature dependence of embryonic development. Lastly, we model coupled chemical reactions to explain this surprising observation.

## Results

### The Arrhenius equation is a good approximation for temperature dependence of embryonic progression

To experimentally investigate the temperature dependence of complex biological systems, we acquired time-lapse movies of fly (*Drosophila melanogaster*) and frog (*Xenopus laevis*) embryos from shortly after fertilization until the onset of movement in carefully controlled temperature environments (Fig. S1). Observing these different embryos allows us to assess the generality of our findings as the species are separated by ∼1 billion years of evolution (Hedges, 2002). Both species’ embryos developed as exotherms and have evolved to be viable over a wide temperature range. Specifically, we find fly embryos are viable in temperatures ranging from ∼14 °C to ∼30 °C, while frog embryos are able to develop from ∼12 °C to ∼29 °C. For fly embryos, we recorded developmental progression over 12 events, which we could reproducibly score from our movies (Fig. 1A, S2A, C, D, Movie S1). Similarly, figure 1B represents the events that we could reproducibly score for frog embryos (Fig. S2B, E, F, Movie S2). Figure 1C shows for each fly embryo stage the mean time since t=0, which we define as the last syncytial cleavage (Fig. S3A). At high temperatures (e.g. 33.5 °C), fly embryos are able to develop until gastrulation, but die at this stage (during germ band shortening). Similarly, for temperatures at and below ∼9.5 °C, fly development arrests after gastrulation (after germ band shortening). We see a general trend of decreasing developmental time for every stage with increasing ambient temperature (Kuntz and Eisen, 2014). Figure 1D shows the developmental times in frog embryos since 3rd Cleavage, which we define as t=0 (Fig. S3B). Here too, we see an inverse relationship between developmental time and temperature (Khokha et al., 2002), suggesting a potential Arrhenius relationship. To investigate how well this temperature dependence can be captured by the Arrhenius equation we obtain pseudo developmental rates by inverting time-intervals between the scored developmental stages. We then generated Arrhenius plots by plotting the natural logarithm of these rates against the inverse of relevant absolute temperatures. If a process strictly follows the Arrhenius’ equation, it appears linear in the Arrhenius plot. Although both the frog and fly data exhibit wide core temperature regions that appear well approximated by a linear fit, between 14.3 and 27 °C in flies and 12.6 and 25.7 °C in frogs (Fig. 2A, B), in each case clear deviations are observed as temperatures near the limits of the viable range and outside the core temperatures (Fig. S3C, D).

**Figure 1:**
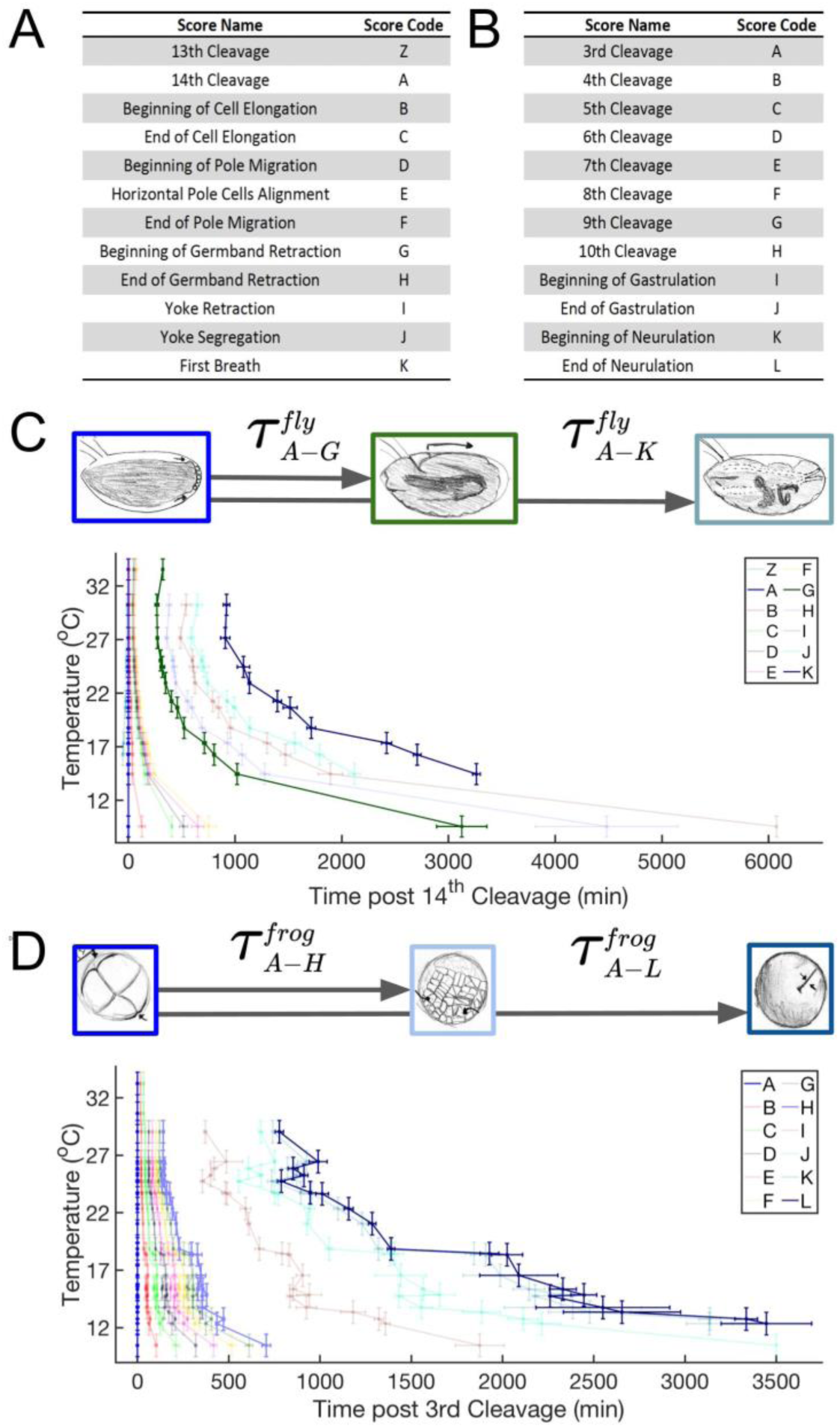
Temperature dependence of development progression in fly and frog embryos. **A)** A table representing *D. melanogaster* developmental scoring event names and sequential stage codes used throughout this paper. **B)** *X. laevis* developmental scoring and codes used throughout this paper. **C)** Shown here is a schematic depicting how time (*τ*) intervals for stages are determined based on beginning and ending score. Plotted also are all time intervals of *D. melanogaster* embryos to reach various developmental scores shown in (A), measured at temperatures ranging from 9.4 °C to 33.4 °C. For each stage we show the time-interval since score A (14^th^ cleavage), which we define as t = 0, up to each scored event. Zoom in of early stages are shown in figure S3A. Shown are the mean of replicates with error bars indicating 95% confidence intervals. **D)** Time intervals of *X. laevis* embryos to reach the various developmental stages measured at temperatures ranging from 10.3 °C to 28.9 °C. For each stage we show the time-interval since stage A (3rd cleavage), which we define as t = 0. Zoom in of early stages are shown in figure S3B. Each interval shows the mean of replicates. Error bars indicate 95% confidence intervals.

**Figure 2:**
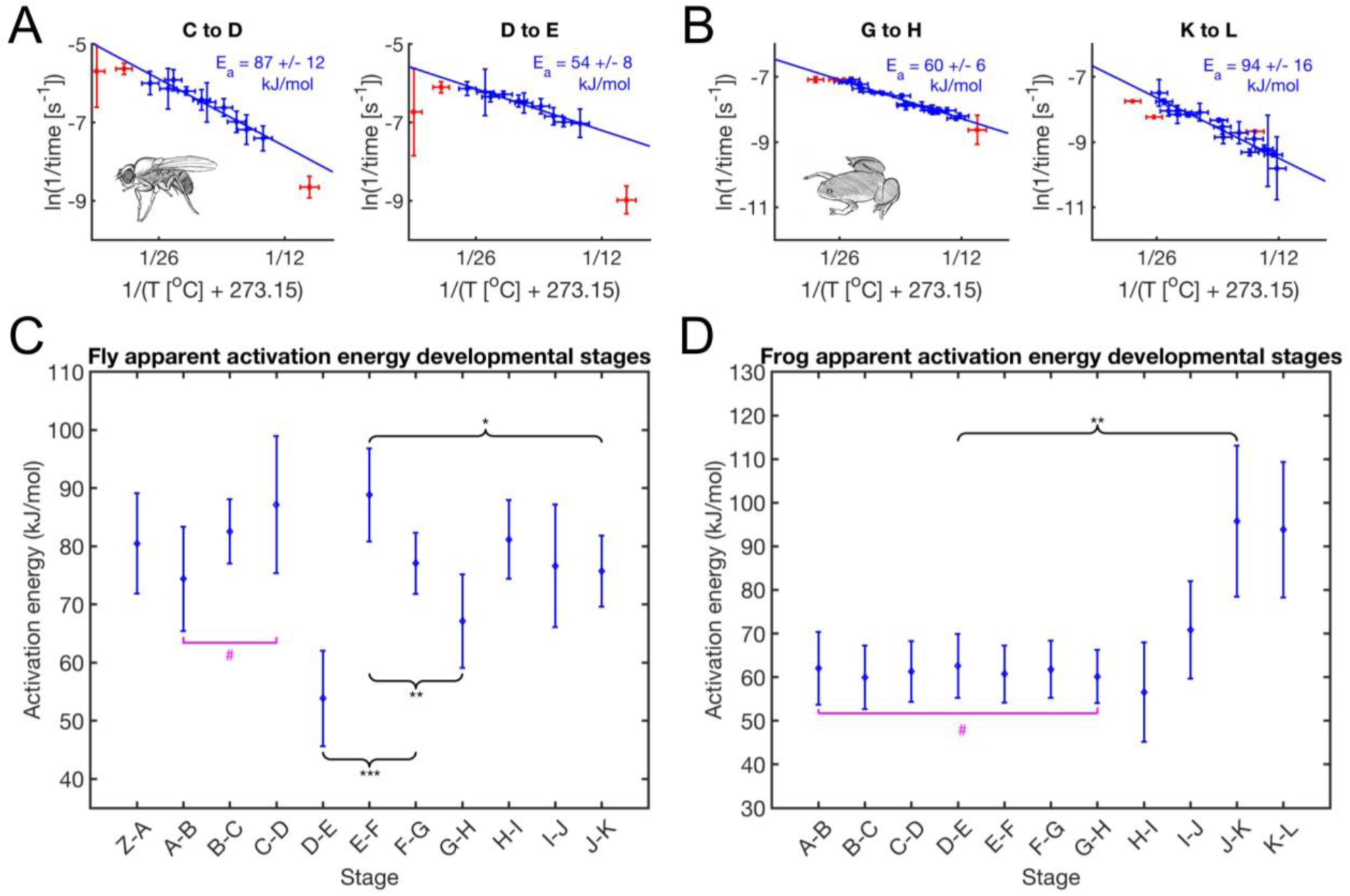
Apparent activation energies vary significantly between developmental stages. **A)** Two examples of developmental time intervals in *D. melanogaster* converted to Arrhenius space. Blue data points represent the core temperature range (14.3 to 27 C) used for the linear fit to determine the apparent activation energy. Red data represents more extreme temperature values with consistently lower than expected rates. **B)** As (A) but for *X. laevis*. Temperature intervals in blue (12.2 to 25.7 C) were used to determine apparent activation energies. **C)** Apparent activation energies for adjacent Fly developmental intervals were calculated from the Arrhenius corresponding Arrhenius plots (Fig S3C). The x-axis is labeled with the endpoint stage of each interval. Each point shows the 95% confidence interval for the activation energy. Magenta brackets represent groupings (all points above the bracket) showing no statistical difference (#) in activation energy. Black braces connect a few examples of stage transitions that did show statistically significant differences in slope (and thus E_a_). *** p < 0.001, ** p < 0.01, * p < 0.05. **D)** As (C) but for frog developmental intervals from plots shown in Fig S3D.

From the well approximated core temeprature ranges we inferred the apparent activation energies from the slope of the Arrhenius plots for all developmental intervals between adjacent scored events. Figure 2A, B shows two example stagings in each *D. melanogaster* and *X. laevis*. From these plots it is apparent that the developmental rates within these stage pairs of each organism show different apparent activation energies (example seen in fly E_a_ = 87 kJ/mol, E_a_ = 54 kJ/mol, p-value of 0.0001, F test (Lomax, 2007)) i.e. their developmental rates scale differently with changing temperatures. In this respect our results differ from the uniform scaling observed for fly development in a previous study (Kuntz and Eisen, 2014), but this difference most likely reflect the greater morphological resolution possible with our imaging setup and the wider temperature range investigated in our experiments. Apparent activation energies for all scored stages in fly embryos range from ∼54 to 89 kJ/mol (Fig. 2C, S4A). This compares similarly to the energy released during hydrolysis of ATP (about 64 kJ/mol) (Wackerhage et al., 1998) and to literature values of enzyme activation energies (∼20-100 kJ/mol) (Lepock, 2005). We performed the equivalent analysis in frog embryos seen in figure 2D. Also here, we observe significantly different activation energies ranging from ∼57 to ∼96 kJ/mol with all cell cycle stages’ activation energies being insignificantly different (E_a_ = 60-63 kJ/mol, p-values between 0.87 and 1, F test) (Fig. S4D). Cell cycle stages are expected to have equivalent Ea’s because of the equivalent processes being performed over each division, therefore consistency over these stages lends some credence to our analysis and application of Arrhenius. Due to the large evolutionary distance between frogs and flies, we cannot compare equivalent stages between these two organisms for most of development. However, the cleavage stages in frog embryos can naively be expected to be driven by similar biochemical events as the syncytial cleavages in fly embryos. Interestingly, the corresponding apparent activation energies are significantly different (p-value = 0.02, F test) between the two species with ∼80 kJ/mol in fly embryos and ∼62 kJ/mol in frog embryos. The cause of this divergence is unknown to us but this may be due to the slight differences between division mechanisms. While different developmental stages show statistically different apparent activation energies this does not interfere with canonical development and the rate of embryogenesis from first to last stages maintains a fairly Arrhenius-like relationship to temperature.

### Measured departures from Arrhenius law

The activation energies calculated for frogs and flies are based on data from the broad core range of temperature that are typically used for lab maintenance of both species. Outside those ranges, the data deviates from idealized behavior and thus is unsuitable for approximating Arrhenius parameters. Figure 3A shows two examples for fly embryos: from the 14^th^ Cleavage to the Beginning of Germband Retraction, and First Breath respectively. We confirmed by Bayesian Information Criterion (BIC) analysis that a quadratic fit is indeed more appropriate than a linear fit over the entire viable temperature range (Fig. 3B) (Dziak et al.; Wit et al., 2012). All quadratic fits over the entire viable temperature range are downward concave (Fig. S3C, D). We performed the identical analyses with all possible developmental intervals between scored events (Fig. 3B). We see a clear statistical preference for a quadratic over linear fit in Arrhenius space for all scored developmental intervals in flies. Figures 3C and 3D show the equivalent analyses performed on frog embryos. Here data is less conclusive as to fit preference, likely due to the overall poorer reproducibility of our frog data but for the majority of scored intervals, including entire development, a quadratic fit is preferred over the linear one (Fig. 3D).

**Figure 3:**
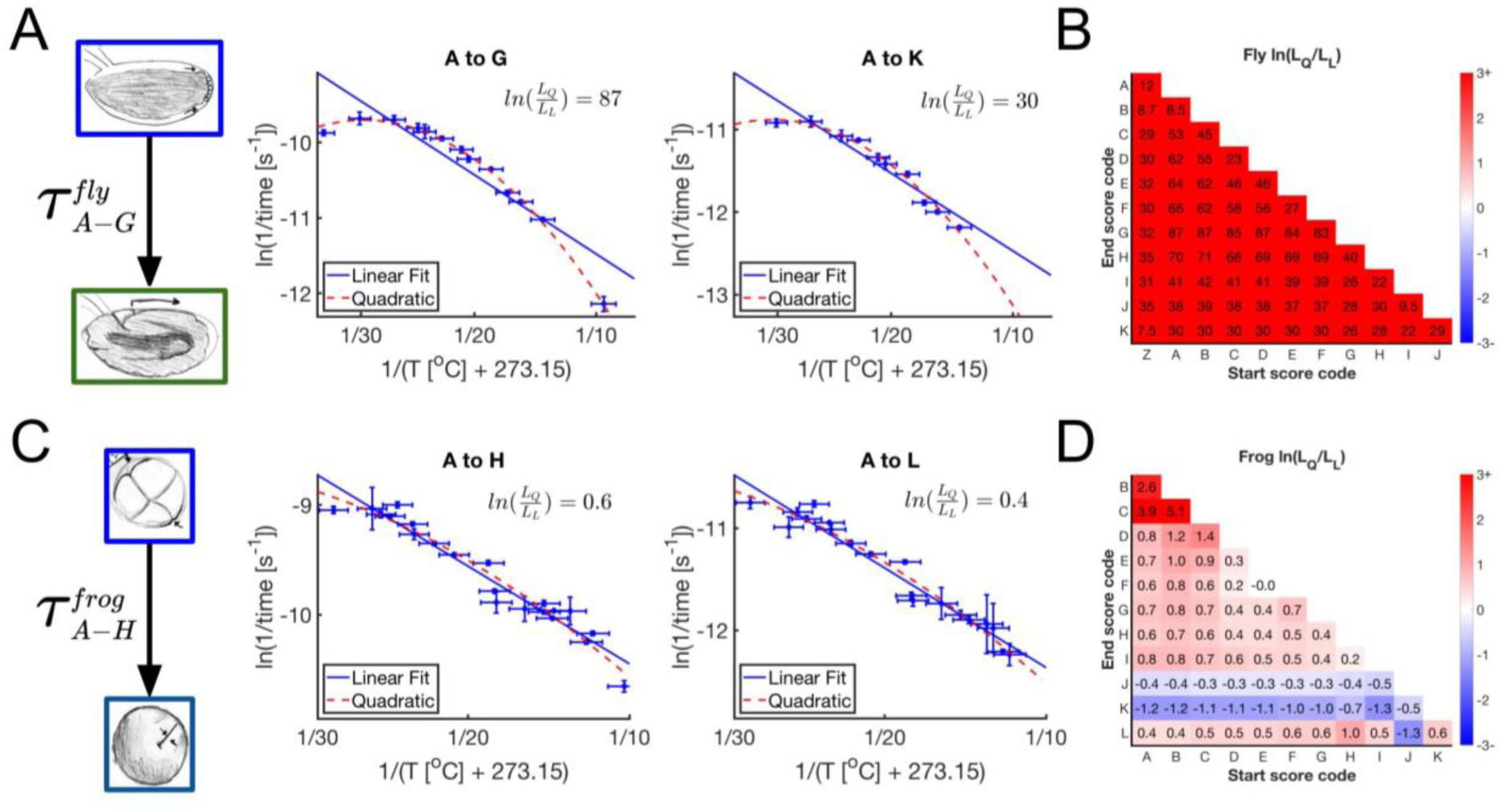
Fitting the temperature dependence of embryonic development with the Arrhenius equation. **A)** Example Arrhenius plots for fly embryos for the interval from last syncytial cleavage (A) to germ band retraction (G) and First Breath (K). At very high and very low temperatures (red), the observed developmental rates are clearly slower than predicted by a linear Arrhenius fit through the core temperature ranges (blue). We can describe the temperature dependence of fly embryonic development over the entire viable temperature range well with a quadratic fit (dashed magenta line). This is confirmed by the log ratio of likelihoods (pink). **B)** The natural log of ratio of BIC (penalized) likelihoods for quadratic over linear. Values above 0 indicate that a quadratic fit is preferred to a linear fit. Here we see such a preference for all scored developmental intervals. **C)** As (A) but for the frog developmental interval from 3rd cleavage (A) to 10th cleavage (H) and End of Neurulation (L). Slight preference is given here to a quadratic fit over linear. **D)** As (B) but for all frog developmental intervals. For most intervals the quadratic model is preferred over the linear model (values above 0). However, this preference is much less clear than in fly and for some intervals the linear fit is preferred.

These findings raise the question if the previous linearly approximated temperature region is also non-linear (14.3 to 27 °C in fly and 12.6 to 25.7 in frog). We performed a Bayesian Information Criterion (BIC) analysis in this narrower range and find that the natural log ratio of the penialized likelihoods for the data of development from stage A to G in fly to be quadratic over linear is ln(L_Quadratic_/L_Linear_) = ∼24 (Fig S4B). We performed equivalent analysis for all developmental intervals in fly embryos and find many instances in which this “linear regime” is actually clearly quadratic (Fig S4C). We always observe deviation to be downward concave i.e. the rates at very low and very high temperatures are lower than predicted by the Arrhenius equation (Fig. S5A). Thus, while the Arrhenius equation is a good approximation for the temperature dependence of early fly and frog development, particularly around the core temperatures, we see clear deviation over both extreme and initially apparent linear temperature ranges. This observation supports our initial intuitions that Arrhenius cannot perfectly describe a complex system, although why it deviates and how it is still a fairly decent approximation remains to be answered.

### Multiple steps and nonideal behavior of individual enzymes lead to non-Arrhenius temperaure dependence

Next, we investigated what could be the cause for the observed non-linear behaviour in the Arrhenius plot for developmental processes (Fig 4A). One possibility is that this observed non-linear behavior arises from the complexity of biological systems composed of multiple coupled elementary reactions, each of which follows Arrhenius. To analyze the conditions under which complex networks consisting of many sequential chemical reactions could be predicted to follow the Arrhenius equation, we analyzed sequential reactions using a relaxation time scale formalism (see supplementary information). For a single reaction we confirmed the known relationship of 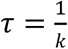. However, when the model is expanded to a relaxation function modeling two reaction transitions, we find equation 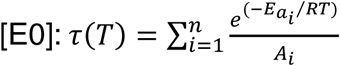, where *τ* is no longer simply related to the inverse of *k*. Converting to Arrhenius coordinates 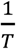 in equation 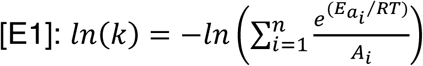, we can clearly see that a sequential multi-reaction series does not yield linear Arrhenius plots. To test whether equation E1 could adequately describe a biological network we investigated how well we could predict the temperature dependence of a large portion of fly’s scored embryonic development based on linear fit parameters for individual developmental events. Interestingly, using parameters extapolated from the linear temperature regime of individual stage intervals we observed that predictions based on this model are a near perfect match to the linear fit for the overall developmental interval (Fig. 4A). We observed the same results for frog embryos (Fig. S6).

**Figure 4:**
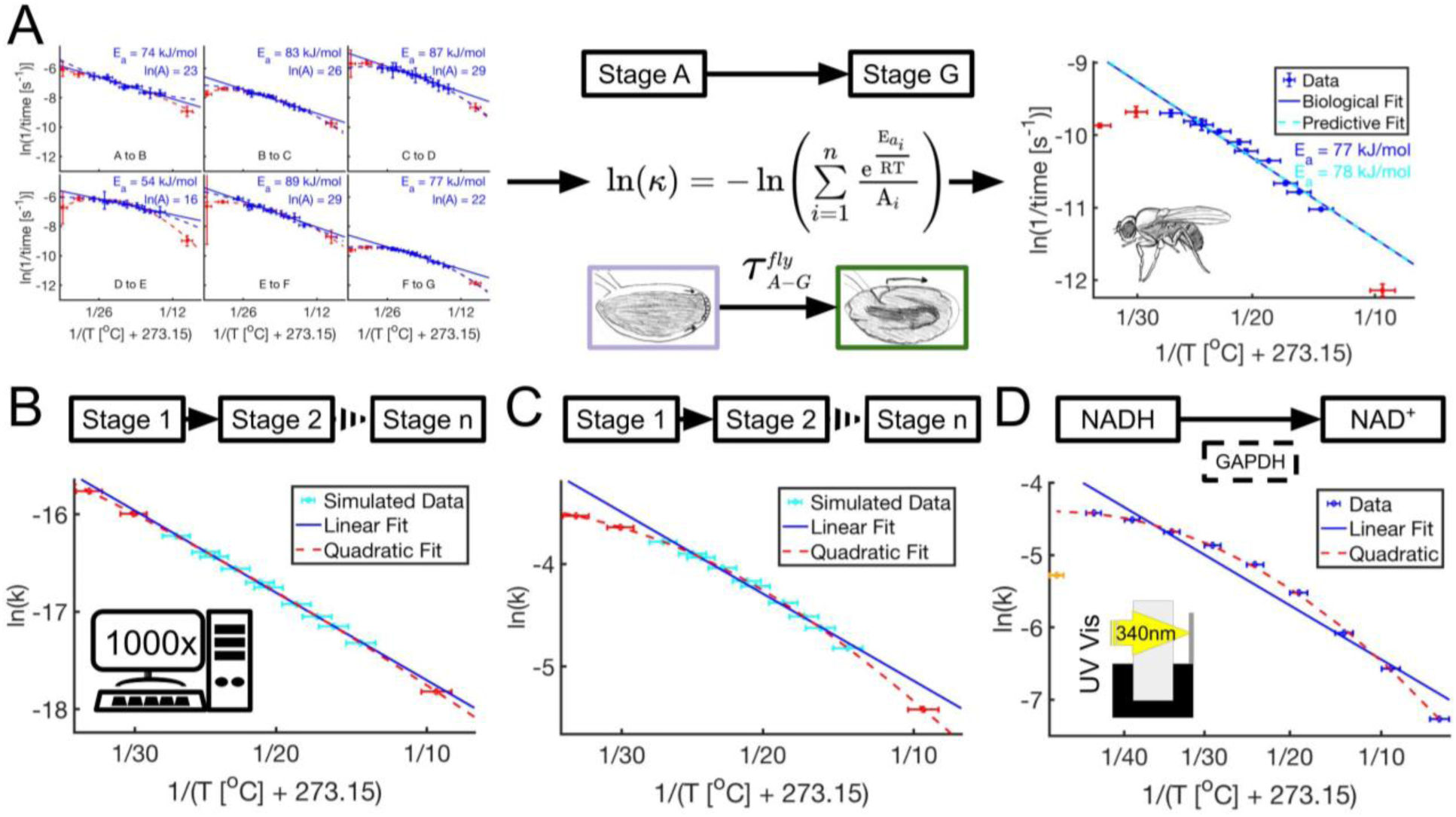
Complexity and non-idealized behavior of individual enzymes can contribute to non-idealized behavior of developmental processes. **A)** Temperature dependence of entire development can be reasonably modeled by coupled sequential Arrhenius processes. We predict the linear regime of entire fly development by coupling the predicted temperature dependence by integrating the measured pre-factors and apparent activation energies for individual stages. This prediction of ln(k) for the composite network shows great agreement with a linear fit of data for the entire developmental interval (A-G). **B)** To investigate whether coupled Arrhenius processes give rise to non-linear behavior in the Arrhenius plot we simulated 1000 coupled reactions with randomly chosen A and E_a_. The entire system ln(k)s were calculated and then fit with a linear fit through the supposed linear temperatures (blue) and a quadratic through the entire temperature range (magenta). We see very minimal deviation from linearity. Error bars indicate 95% confidence intervals for temperature based on our experimental setup. **C)** As (B) however instead of simulating a random sequence of reactions we optimized the chosen A and E_a_ to maximize the curvature at T = 295 °K (dashed magenta). Modest divergence from linearity is observed over the measured temperature range. However, even in this extreme case the observed curvature is less than our biologically observed (A). **D)** To investigate whether non-linear behavior of individual enzymes could contribute to non-idealized behavior of complex biological systems, we investigated the temperature dependence of the activity of GAPDH. GAPDH’s conversion of NADH to NAD+ can be conveniently monitored with UV/VIS spectroscopy at 340nm. Interestingly, this enzyme shows clearly non-idealized behavior from 9 to 34 °C (vertical gray lines) and beyond, comparable to what we see for embryogenesis as a whole. At very high temperature the rates are actually decreasing with increased temperatures likely due to protein denaturation (orange).

We wondered if we could generalize this observation and show that even though coupled Arrhenius equations are technically non-linear, they might appear linear under biological experimentally accessible scenarios. To this end, we used our equation (E1) to simulate many (1000) sequentially coupled chemical reactions, each defined by its own activation energy and prefactor and observed how the reaction rates of the entire network scale with temperature. When we randomly choose 1000 A’s and Ea’s among reasonable ranges, we observe nearly perfect linear behavior in the Arrhenius plot for the entire system (Fig. 4B, S5B). Next, we optimized E_a_ and A to maximize the curvature of the system at T = 295 K, while constraining E_a_ between literature values 20-100 kJ (Lepock, 2005) and constraining the time 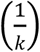 for embryonic states between 1 second and 3 days. We chose 1 second as lower limit as an unreasonably short time in which an embryo could transition through a distinct biochemical state. We used the 3 days of entire fly development in our experiment as an upper time limit for distinct embryonic states. When maximizing curvature for 1000 coupled reactions using these limits, we observed some non-linearity, which we believe could be experimentally detectable, even over the core temperature range (Fig. 4C, S5C). Analysis of our equation E1 shows consistant downward concave divergence from linearity (Fig. S5A). However, even assuming these “worst case” scenarios, our simulations do not show the same qualitative level of divergence from linearity observed in our biological data in figure 4A. Furthermore, in our data we observe a decrease in reaction rates at very high temperatures (Fig. 4A). This is impossible to achieve with simulations of coupled reactions that follow the Arrhenius equation. This suggests that factors other than coupling of many reactions are likely to contribute to the non-linearity in the Arrhenius plot of the temperature dependence of developmental processes.

Is it possible that the individual steps, e.g. enzymes, are already non-Arrhenius? To investigate this possibility, we measured the temperature dependence of a reaction catalyzed by the model enzyme GAPDH, a glycolytic enzyme. We chose GAPDH because it is essential to all forms of eukaryotic life and its activity can be easily assayed by following the increase of absorbance of NADH/NAD+ at 340 nm with spectrophotometry. Interestingly, we find that GAPDH shows clearly non-linear behavior in the Arrhenius plot from 4 to 44 °C (Fig 4D). As with developmental data we find that GAPDH activity follows concave downward behavior. Similar to our embryonic data, we find that reaction rates at the high end reduce with increasing temperature (Fig. 4A, D). At the very high end the enzyme is likely starting to denature (Daniel et al., 1996). However, denaturing is unlikely to explain the non-idealized behavior at lower temperatures. It has been proposed that such downward concave behavior in the Arrhenius plot could be due to changes of rate-limiting steps as a function of temperature (Fersht, 1999) or due to lower heat-capacity of the transition state versus enzyme substrate complex (Arcus and Mulholland, 2020; Arcus et al., 2016; Hobbs et al., 2013). The enzyme might have evolved to work optimally at a certain temperature (39 °C for rabbit GAPDH used in this assay). Deviating to lower or higher temperatures might lead to reaction rates that are lower than when assuming idealized Arrhenius behavior. Consistent with this, all Arrhenius plots shown for developmental stages or this individual enzyme are statistically significant quadratic with downward concave shape (Fig. 3B, S5A).

## Discussion

The Arrhenius equation is used for simple chemical reactions to relate the reaction rate with the energy necessary to overcome activation barriers, i.e. activation energy. We have shown that embryonic fly and frog development can be well approximated by the Arrhenius equation over each organism’s core viable temperature range. Using this relationship, we can show that the apparent activation energies for different developmental stages can vary significantly, showing that different parts of embryonic development are scaling differently with varying temperatures. By examining the data more carefully, we observe that the relationship between temperature and developmental rates in both species is confidently better described by a concave downward quadratic in Arrhenius space, especially when considering the entire temperature range over which the embryos are viable. Our modeling studies demonstrate that in principle, linking multiple Arrhenius governed reactions could lead to concave downward behavior. However, when we reasonably limit k and E_a_, coupling of sequential reactions can only modestly contribute towards the observed non-idealized behavior. In contrast, when we observed the temperature dependence of a single enzymatic model process, the assay of GAPDH, we observed clear non-linear, concave downward behavior. Therefore, the observed curvature of developmental rates can be most easily explained by individual non-idealized rate-limiting processes.

One striking finding of our study is that different developmental processes within the same embryo clearly scale differently with varying temperature. We observe this not only in the different apparent activation energies at different stages but also in the different divergence from non-linearity at extreme temperatures. One major question that still remains is how complex embryonic development can result in a canonical developed embryo if the different reactions required for faithful development proceed at different relative speeds at different temperatures. In our assays we are only able to follow temporally sequential reactions, and one can argue that increasing or decreasing time spent at a particular stage should not influence the success of development. However, development must be much more complex and hundreds or thousands of reactions and processes must occur in parallel, e.g. in different cell types developing at the same stage. How can frog and fly embryos be viable over ∼15 °K temperature range over which stages’ different temperature sensitivity could possibly throw development out of balance? We see two possible developmental strategies to overcome this problem. Either all rate-limiting steps occurring in parallel at a given embryonic stage have evolved similar activation energies, or the embryos have developed checkpoints that assure a resynchronization of converging developmental processes over wide temperature ranges.

## Acknowledgement

We would like to thank Trudi Schüpbach and members of the Wühr and Wieschaus labs for helpful suggestions and discussions. This work was supported by NIH grant R35 GM128813 (MW), R01 GM134204-01 (SS) and T32 GM007388 (NP). We are grateful for HHMI support (EW).

