## Supplementary Information for "Evaluating the Simple Arrhenius Equation for the Temperature Dependence of Complex Developmental Processes"

### Supplementary Figures

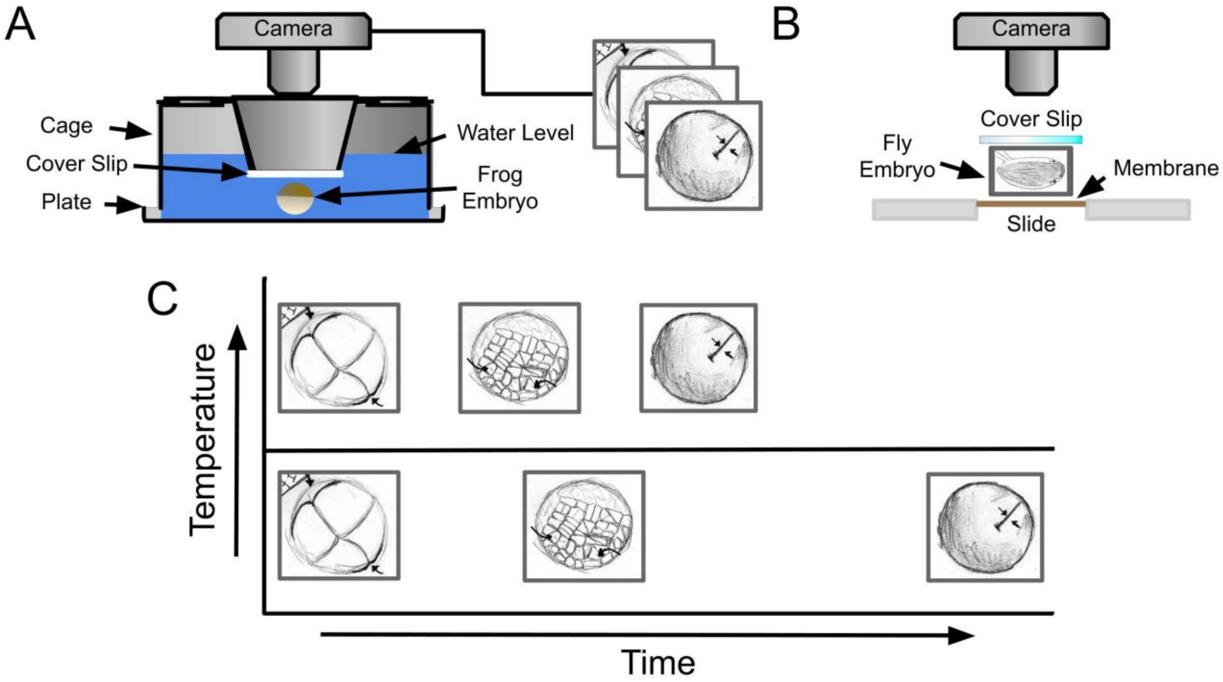


**Supplementary Figure 1: Process for collecting and organizing developmental data. A)** A schematic of the process used to collect frog developmental data. Frog embryos were fertilized, de-jellied, and then placed on a custom water filled plate. The plate was then covered with a custom cage with a conical recess to control water depth above the embryo. 50 mL of acclimated water was then poured in through the holes in the cage roof. Placed under a camera attached to a microscope, the embryos were allowed to develop while being recorded; connecting line shows output for producing images shown to the right. **B)** Shown here is the process for setting up fly embryos in Halocarbon oil on an air permeable slide for time-lapse imaging. **C)** Embryos were recorded at various temperatures. Various milestones were scored and graphed against their temperature as shown here.

A

| Cut Score Name | Cut Score Code |
|---|---|
| Beginning of PMG Opening | 1 |
| Head Invagination | 2 |
| Gut Movement | 3 |
| Head Movement | 4 |
| End of First Breath | 5 |
| Gut Contents Migration | 6 |
| Hatching | 7 |

B

| Cut Score Name | Cut Score Code |
|---|---|
| Formation of Optic Lines | 1 |
| De-Sphericalization | 2 |
| Head Becomes Distinct | 3 |
| First Muscle Twitch | 4 |

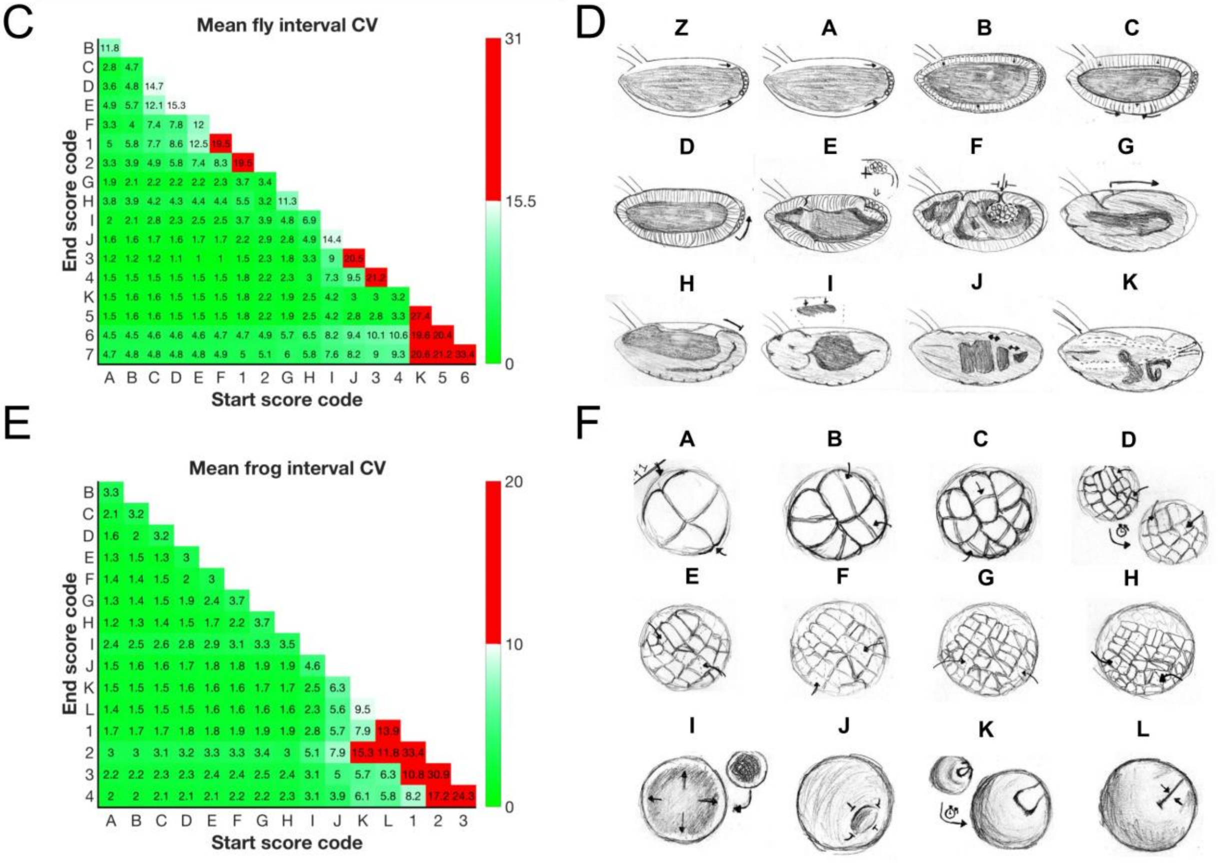


**Supplementary Figure 2: Animal developmental stages and coefficient of variance (CV) analysis. A)** Seven additional developmental events (scores) in fly that were later cut. Given is a descriptive name and number score code for reference in the following CV analysis. **B)** As (A) but for four additional cut frog developmental events. **C)** CV's were calculated for every developmental time interval between scores we considered investigating and for every temperature. Means were calculated over all temperatures for each interval to represent variation for each developmental interval. CV's are displayed as percentages. Color is used to differentiate the CV cutoff we used to distinguish which 12 stages were reproducible enough to reliably score, red (15.5) being above the cutoff. **D)** 12 developmental stages with CVs below 15.5, determined to be the most reproducible, in *D. melanogaster*. **E)** As (C) but for 12 frog developmental scores. **F)** 12 developmental stages we investigated in *X. laevis* determined most reproducible with CVs below 10.

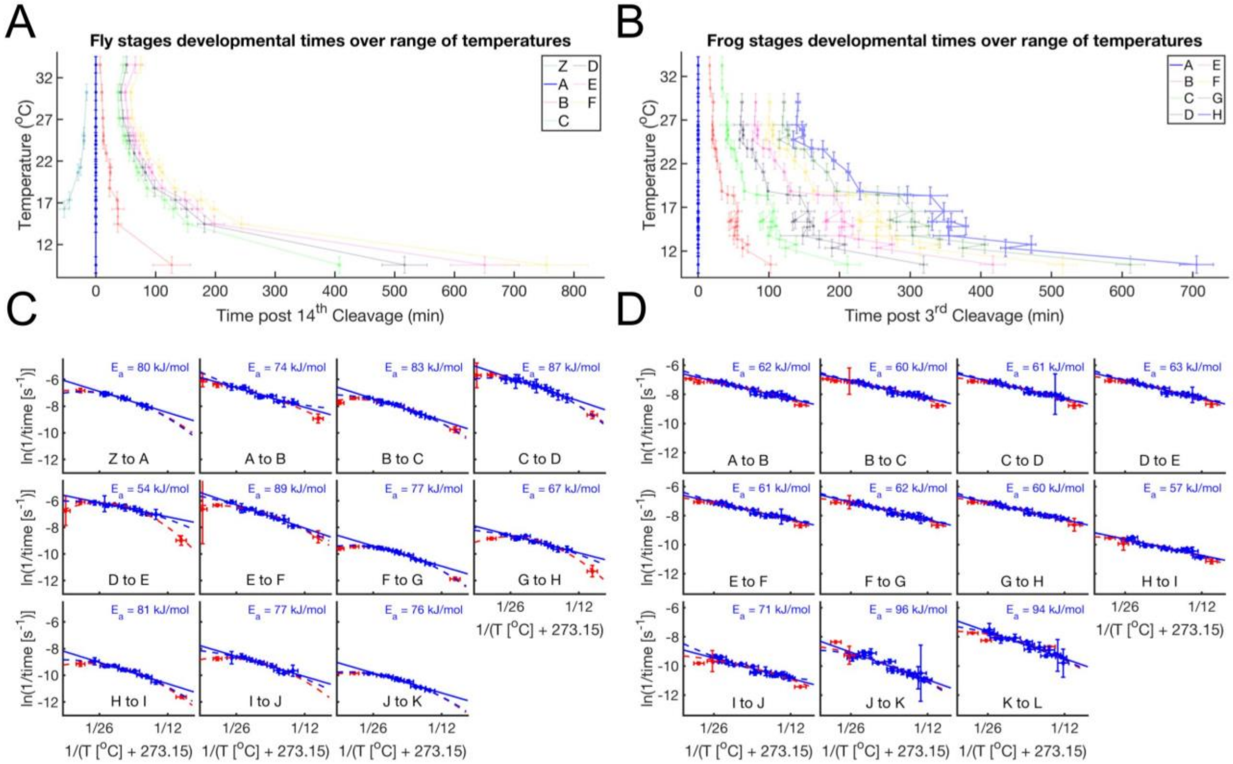


**Supplemental Figure 3: Different temperature dependence of developmental progression in frog and fly embryos. A)** As Figure 1B but zoomed in on the relatively rapid early fly developmental stages. **B)** As Figure 1D but zoomed in on the relatively rapid frog cleavage stages. **C)** Shown are Arrhenius Plots (similar to Fig 2 A) for 10 adjacent fly developmental stages. Linear fits were calculated only on the blue data points (over the range previously shown in Fig 2A). The apparent activation energy for each interval is displayed top right (in blue) of every subplot. Apparent activation energies were calculated by multiplying the slope by -R (in kJ). Clearly shown here is the consistency for extremes (red data) to fall below the expected Arrhenius linear fit prediction, but are captured by the magenta quadratic fit through all the data, including extremes. Additionally the dashed blue line shows a quadratic fit for the interior blue data points as a visual guide for its true non-linear behavior. **D)** As (C) but using 11 adjacent frog developmental intervals and calculated over the interior temperature range used in Figure 2B.

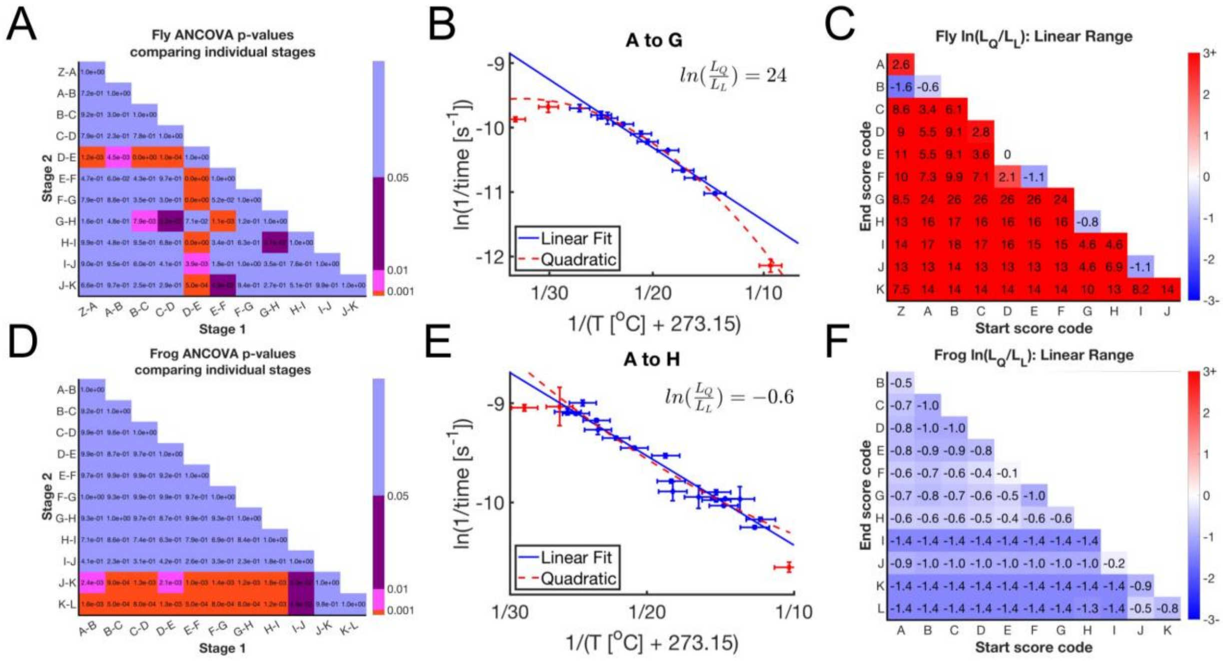


**Supplemental Figure 4: Statistics, p-values and BIC comparisons over the linear range.** **A)** Fly p-values using F-test (ANCOVA) calculated to determine statistically significant difference between every combination of linear fit slopes from developmental intervals shown in Fig. S3C. As in Fig. S3C Stage 1 and 2 are marking the event by which then end of the stage is scored since the last event. Since A is T0 it is left blank while we calculate backwards to Z ($13^{th}$ cleavage). Blue represents p-values above 0.05, purple is ≤0.05, pink is ≤0.01, and red is ≤0.001. **B)** Linear and quadratic fits were drawn over the interior temperature range (blue data, excluding extreme red data) of fly development. Data still shows a strong preference for quadratic fits (magenta) with a value about 24 for log ratio likelihood of quadratic over linear. **C)** The natural log of BIC comparison between quadratic and linear (log ratio likelihood of quadratic over linear) are shown here for all developmental intervals in fly development over the interior temperature range. Intervals are marked with their beginning event on the X-axis, and their ending event on the Y-axis. Here we see the majority of intervals still strongly prefer a quadratic fit (magenta) over a linear fit (blue). Even where a linear fit is preferred it is very weakly preferred (near zero). **D)** As (A) but for calculating frog p-values between slopes of developmental intervals shown in Fig. S3D. **E)** As (B) but for frog 3rd to 10th cleavage and showing slight, but weak, preference for linearity. **F)** As (C) but for all frog intervals. Here we see very slight preference for a linear fit, which may be due to noise cause by maternal batch effects and feeding inconsistency that are likely not present in fly.

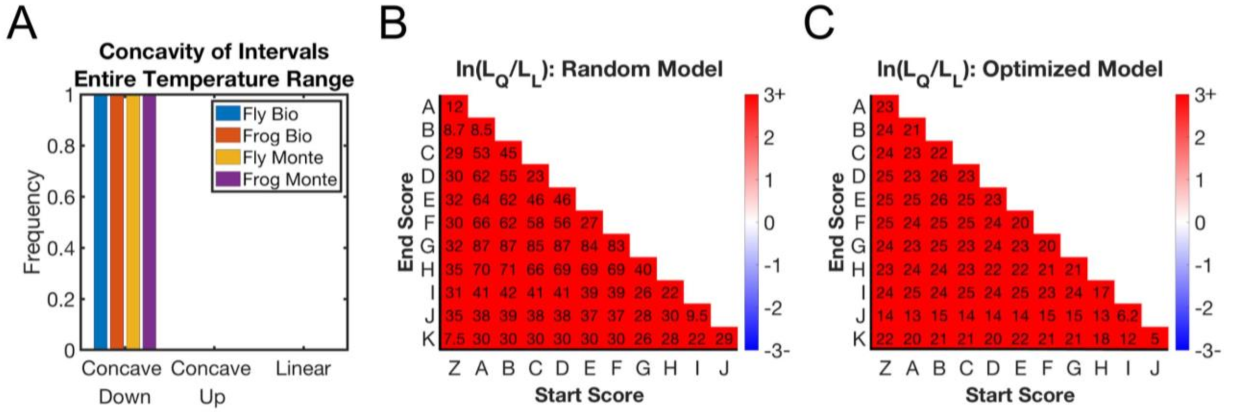


**Supplementary Figure 5: Concavity of developmental intervals and Monte Carlo simulations of Fig. 4C&D show model ability to contribute to nonlinearity. A)** Shown here is a comparison of the frequency of concave downward Arrhenius plots for our biological data (over every interval over entire temperature range) and Monte Carlo simulations using biological errors in time and temperature (entire range) for every developmental interval with Fig. 4C as the assumed true underlying model. Each developmental interval had 100x Monte Carlo simulations performed. **B)** A heatmap showing the natural log ratio of penalized likelihoods (quadratic/linear) comparing fit preference for every developmental interval in fly using that interval's standard error in time and temperature (14.3 to 27 C) with Fig. 4B as the true underlying model. Each interval shown is the median of 100x simulations. **C)** As (B) but using Fig. 4C as the true underlying model.

A

| Stage | Fly lnA | Fly Ea |
|---|---|---|
| Z to A | 25 | 80 |
| A to B | 23 | 74 |
| B to C | 26 | 83 |
| C to D | 29 | 87 |
| D to E | 16 | 54 |
| E to F | 29 | 89 |
| F to G | 22 | 77 |
| G to H | 18 | 67 |
| H to I | 24 | 81 |
| I to J | 22 | 77 |
| J to K | 21 | 76 |

B

| Stage | Frog lnA | Frog Ea |
|---|---|---|
| A to B | 18 | 62 |
| B to C | 17 | 60 |
| C to D | 18 | 61 |
| D to E | 18 | 63 |
| E to F | 17 | 61 |
| F to G | 18 | 62 |
| G to F | 17 | 60 |
| F to I | 13 | 57 |
| I to J | 19 | 71 |
| J to K | 29 | 96 |
| K to L | 30 | 94 |

C

$$\ln(\kappa) = -\ln\left(\sum_{i=1}^{n} \frac{e^{\frac{E_{a_i}}{RT}}}{A_i}\right)$$

D

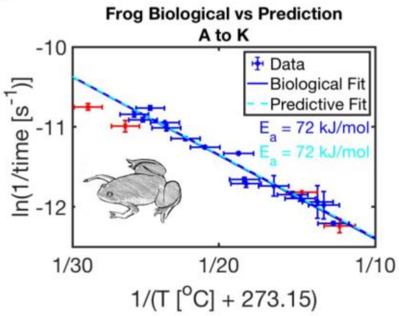


**Supplemental Figure 6: Parameters for linear fits and sequential linear model prediction capture temperature dependence of entire embryonic development well. A)** A table showing fly stages marked by their start and ending events and their empirically determined prefactor (lnA) and apparent activation energy ($E_a$) from Fig. S3C. **B)** As (A) but for frog prefactor and apparent activation energy calculated empirically from Fig. S3D. **C)** Our predictive equation determined in E1 for an assumed sequential linear network. **D)** A prediction (dashed cyan) for the Arrhenius plot of frog developmental interval A (3rd cleavage) to K (Beginning of Neurulation) with the first 10 parameters pairs given in (B) and the equation in C. The prediction is plotted against the biological data and linear fit (solid blue) over the temperature interval 12.2 to 25.7 $^{o}$C. Here we see a near perfect agreement. Stages were selected to conserve unique temperature data points, as missing or inconsistent data skews the prediction.

### Supplementary information
#### Math derivations

*Multi-reaction Network*

The following is a derivation for an equation modeling the temperature dependence of a sequential multi-reaction system of reactions, each of which individually follow Arrhenius.

First we show that a relaxation time scale formulation can adequately yield the known $\tau = \frac{1}{k}$, assuming a simple transition from some stage $A \rightarrow B$.

$$R_{A\rightarrow B}(t) = \frac{A_0 - B(t)}{A_0}$$

$$R_{A\rightarrow B}(t) - R(t+\tau) = -\frac{\partial R}{\partial t}dt$$

$$\tau = \int_0^\infty t\left(-\frac{\partial R}{\partial t}\right)dt$$

$$= tR(t)|_0^\infty + \int_0^\infty R(t)\,dt$$

$$= \int_0^\infty R(t)\,dt$$

$$R(t) = \frac{A_0 - B(t)}{A_0}$$

Here we substitute $A_0 - A_0e^{-kt}$ for $B(t)$

$$R(t) = \frac{A_0 - (A_0 - A_0e^{-kt})}{A_0}$$

$$= e^{-kt}$$

Now we substitute $e^{-kt}$ for $R(t)$ in $\tau = \int_0^\infty R(t)dt$

$$= \int_0^\infty e^{-kt}dt$$

$$= \frac{1}{k}e^{-kt}|_0^\infty$$

$$\tau = \frac{1}{k}$$

Therefore, for a simple transition from A to B a relaxation formulism captures the known relationship of $\tau = \frac{1}{k}$ and by substituting in the Arrhenius equation $\tau = \frac{e^{Ea/RT}}{A}$.

Now we investigate a more complicated transition series, from $A \rightarrow B \rightarrow C$ and determine if a relaxation formulism can be generalizable to arbitrarily high reaction transitions or stages.

$$R(t) = \frac{A_0 - C(t)}{A_0}$$
$$A(t) = A_0 e^{-k_1 t}$$
$$B(t) = A_0 \frac{k_1}{k_2 - k_1}(e^{-k_1 t} - e^{-k_2 t})$$
$$C(t) = A_0 \frac{k_1}{k_2 - k_1}\left(k_2(1 - e^{-k_1 t}) - k_1(1 - e^{-k_2 t})\right)$$
$$R(t) = \frac{A_0 - A_0 \frac{k_1}{k_2 - k_1}\left(k_2(1 - e^{-k_1 t}) - k_1(1 - e^{-k_2 t})\right)}{A_0}$$
$$= 1 - \frac{1}{k_2 - k_1}\left(k_2(1 - e^{-k_1 t}) - k_1(1 - e^{-k_2 t})\right)$$
$$= \frac{(k_2 - k_1) - (k_2 - k_2 e^{-k_1 t} - k_1 + k_1 e^{-k_2 t})}{k_2 - k_1}$$
$$= \frac{k_2 - k_1 - k_2 + k_2 e^{-k_1 t} + k_1 - k_1 e^{-k_2 t}}{k_2 - k_1}$$
$$= \frac{k_2 e^{-k_1 t} - k_1 e^{-k_2 t}}{k_2 - k_1}$$

We can now substitute $R(t)$ back into $\tau = \int_0^\infty R(t)dt$

$$= \frac{1}{k_2 - k_1}\int_0^\infty (k_2 e^{-k_1 t} - k_1 e^{-k_2 t})dt$$
$$= \frac{1}{k_2 - k_1}\left(k_2 \int_0^\infty e^{-k_1 t}\, dt - k_1 \int_0^\infty e^{-k_2 t}\, dt\right)$$
$$= \frac{1}{k_2 - k_1}\left(k_2\left(\frac{1}{k_1} e^{-k_1 t}\Big|_0^\infty\right) - k_1\left(\frac{1}{k_2} e^{-k_2 t}\Big|_0^\infty\right)\right)$$
$$= \frac{\frac{k_2}{k_1} - \frac{k_1}{k_2}}{k_2 - k_1}$$
$$= \frac{\frac{k_2^2 - k_1^2}{k_1 k_2}}{k_2 - k_1}$$
$$= \frac{(k_2 - k_1)(k_2 + k_1)}{(k_1 k_2)(k_2 - k_1)}$$
$$= \frac{k_2 + k_1}{k_1 k_2}$$
$$= \frac{1}{k_2} + \frac{1}{k_1}$$

We can now substitute in the Arrhenius equation, $k = Ae^{-E_a/RT}$, for $k$ to give us $\tau$ in terms of Arrhenius parameters $A$ and $E_a$.

$$\tau(T) = \frac{e^{E_{a_2}/RT}}{A_2} + \frac{e^{E_{a_1}/RT}}{A_1}$$

Here we see a pattern, and further analysis (not shown) of higher order reactions series confirms its generality. The above equation can be simplified to:

$$E0: \tau_n(T) = \sum_{i=1}^{n} \frac{e^{E_{a_i}/RT}}{A_i}$$

Now, since we know that $\tau = \frac{1}{k}$, we can put E0 in terms of $k$ instead of $\tau$ and arrive at a form close to the linearized Arrhenius equation form, $ln(k) = ln(A) - \frac{E_a}{RT}$.

$$k(T) = \tau(T)^{-1}$$

$$= \left(\sum_{i=1}^{n} \frac{e^{E_{a_i}/RT}}{A_i}\right)^{-1}$$

$$ln(k) = ln\left(\left(\sum_{i=1}^{n} \frac{e^{E_{a_i}/RT}}{A_i}\right)^{-1}\right)$$

$$= -ln\left(\sum_{i=1}^{n} \frac{e^{E_{a_i}/RT}}{A_i}\right)$$

Therefore, for a sequential series of $n$ reactions, each of which strictly follows Arrhenius, then the Arrhenius plot for such a system should follow the below equation.

$$E1: ln(k) = -ln\left(\sum_{i=1}^{n} \frac{e^{E_{a_i}/RT}}{A_i}\right)$$

*Concavity*

The following is a derivation for the concavity of E1, which models the temperature dependence of a sequential multi-reaction system of reactions, each of which individually follow Arrhenius.

We begin with equation E1: $\ln(k) = -\ln\left(\sum_i \frac{e^{E_{a_i}/RT}}{A_i}\right)$. To simplify calculations, let $x_i = \frac{e^{E_{a_i}/RT}}{A_i}$, and let $c_i = E_{a_i}/R$. Then, $\frac{dx_i}{d(1/T)} = c_i x_i$.

Now, if we let $S = \sum_i \frac{e^{E_{a_i}/RT}}{A_i} = \sum_i x_i$, then $S' = \frac{dS}{d(1/T)} = \sum_i c_i\, x_i$ and $S'' = \frac{dS}{d(1/T)} = \sum_i c_i^2\, x_i$.

Thus, we have

$$\frac{d(\ln(k))}{d(1/T)} = \frac{d}{d(1/T)}(-\ln(S)) = -\frac{S'}{S},$$

and then

$$\frac{d^2(\ln(k))}{d(1/T)^2} = -\frac{SS'' - (S')^2}{S^2}.$$

We claim that this second derivative is always negative. Since the denominator is a square and thus positive, it is sufficient to show that $SS'' - (S')^2 > 0$. But we know that

$$SS'' - (S')^2 = \left(\sum_i x_i\right)\left(\sum_i c_i^2 x_i\right) - \left(\sum_i c_i x_i\right)^2.$$

Now, consider the coefficient of $x_i x_j$ in this difference. If $i = j$, then both terms will give a coefficient of $c_i^2$, and these will cancel each other out in the difference. Otherwise, if $i \neq j$, then the first term will give a coefficient of $(c_i^2 + c_j^2)$, while the second term will give a coefficient of $2c_i c_j$. Therefore, the resulting term in the difference of these products will be

$$(c_i^2 + c_j^2 - 2c_i c_j)x_i x_j = (c_i - c_j)^2 x_i x_j.$$

Since each $x_i$ is positive (being the quotient of an exponential, which is always positive, and a positive prefactor $A_i$), and since $(c_i - c_j)^2$ is a square, the entire term $(c_i - c_j)^2 x_i x_j$ is positive (unless $c_i = c_j$, in which case it's equal to $0$). Therefore, unless all the $c_i$'s, and thus all the $E_{a_i}$'s, are equal, we have that $SS'' - (S')^2 > 0$, and so the second derivative $\frac{d^2(\ln(k))}{d(1/T)^2}$ is always negative. This means that the graph of $\ln(k)$ versus $1/T$ is always concave down.

#### Materials and Methods:

##### Drosophila melanogaster data collection and analysis

*D. melanogaster* females mutant for Klarsicht were maintained as previously described (Jäckle and Reinhard, 1998; Wieschaus and Nüsslein-Volhard, 1986). Flies were allowed to lay eggs for an hour, at which time fresh embryos were collected for time lapses. Embryos were submerged in halocarbon oil 27 (Sigma Cat# H-8773) and selected if they were retracted from the posterior vitelline membrane, signaling successful fertilization.

3 - 4 selected fly embryos were mounted in halocarbon oil on a slide with an oxygen permeable membrane (Kenneth Technology, Biofoil #03-670-814). A glass coverslip was gently rolled over the embryos. Slides were placed on transmitted light bright field microscopes set at 20x magnification in temperature controlled rooms for between 9.4 and 33.4 C. Images were acquired with one of the following cameras: Canon Rebel Ti5/6, Swiftcam 3 Megapixel, OMAX 9.0 MP, AmScope MU300 and associated software. Time lapses were recorded from syncytial cleavages until embryo hatching. For data analysis, time lapse images were converted into video files and manually scored based on the scoring metric depicted in Fig. S2. Videos will be available at the ASCB image library upon publication of this work: https://www.ascb.org/science-news/the-cell-an-image-library/

Videos were scored based on video absolute time. Stages times were then calculated relative to 14th cleavage (time zero). To adjust these times the 14th cleavage absolute time was subtracted from all subsequent (or previous) score times.

**Xenopus embryo data collection and analysis**

X. laevis egg and testis were collected according to previous protocols with the following modifications (Wlizla et al., 2018). Female X. laevis frog were induced using 500ul of 1000u HCG (Human chorionic gonadotropin) about 16 hours before egg collection. Females were gently squeezed and eggs harvested dry on a petri dish. A male frog was euthanized in 0.1% aminobenzoic acid ethyl ester (Tricaine, MS222) (Sigma A-5040)) and then sacrificed by pithing. Testes were collected and stored in a 2 mL eppendorf tube with 1x MMR (Ubbels et al.). Later testes were transferred to an oocyte culture media (1 L of OCM; 1 bag of Leibovitz's L-15 Medium powder (ThermoFisher Scientific #41300039), 8.3 mL Penn/Strep, 0.67 g BSA) with pH adjusted to 7.7 by Na+ and filtered through a 0.22 um filter (Mir and Heasman, 2008).
Fertilized and de-jellied eggs were prepared as previously described with minor modifications (Ubbels et al.). About one quarter of one testis, collected from a male frog, was crushed with a pestle and mixed into 400 ul of 1x Marc's Modified Ringer's, 0.1 M NaCl, 2.0 mM KCl, 1 mM $MgSO_4$, 2 mM $CaCl_2$, 5 mM HEPES, 0.1 mM EDTA (Ubbels et al.) to prepare a source of parental sperm. This aliquot was pipetted over about 200 eggs. Using a sterile pestle the solution and eggs were mixed and incubated at room temperature for 5 minutes. Embryos were agitated a second time and incubated for another 5 minutes. Fertilization was then induced using MiliQ $H_2O$. Briefly embryos were then de-jellied. 50 ml a 2% Cysteine de-jellying solution was prepared and its pH titrated to 7.8 with NaOH. MiliQ $H_2O$ in the fertilized embryo dish was then exchanged with the de-jellying solution. Embryos were soaked in this solution for 5 minutes or until the jelly coats appeared to be separated from the embryos. Embryos were then washed three times with MiliQ $H_2O$. Washes were then repeated with 0.1x MMR, and allowed to rest in the final 0.1x MMR wash.
Fertilized and de-jellied embryos were viewed and selected using bright field microscopy for synchronous embryos entering NF stage 2 (first cleavage event) (Nieuwkoop and Faber, 1994). Using a custom made wire-loop pipette the embryos were gently and quickly segregated. Stage 2 embryos were then gently transferred via large pipette to a new petri dish containing temperature equilibrated 0.1x MMR.
A 3D printed cover was created using the OnShape.com online software as an .stl file, which was then 3D printed. The stl file is available on github: https://github.com/wuhrlab/Xenopus-Petri-Cage-Top. A simplified side view can be seen in figure S1A. A special embryo cage was then constructed using the above cover, a petri dish with 5mm mesh fused to the, and clay around the edges (Fig. S1A).
An eyepiece camera attached to a bright field microscope captured a developmental time-lapse for a specific temperature setting. 75ml of 0.1x MMR was equilibrated, at temperatures between 10.3 and 33.1 $^{o}$C, in the incubator several hours before embryo incubation. The embryo plate was placed in the incubator on a bright field microscope-camera stage beneath the objective and a LED ring lamp. The plate was filled with 25mL of 0.1x MMR. Stage 2 embryos were then transferred to the plate and gently

positioned on the mesh. The 3D printed cage was then placed over the embryos. The remaining 50ml 0.1x MMR was pipetted into the cage through the top ventilation holes. A temperature recorder (Elitech RC-5) was placed near the embryo. Temperature recording was initiated shortly before the incubator was closed and the time-lapse initiated. A time-lapse was then taken using a consumer camera or eyepiece camera attached to a bright field microscope at 30-second to 1-minute intervals for all time courses, until stage NF 34, which immediately precedes hatching. All our movies will be uploaded upon publication.

#### Generating Arrhenius Plots for embryonic time courses and subsequent analysis

A pseudo reaction rate was determined by inversion of the interval times. The natural log of this pseudo reaction rate was then plotted against the inverse of absolute temperature (in Kelvin). A linear regression was taken using the means of temperature sets over the interval of temperatures that appear most linear (14.3 to 27 $^{o}$C). The 95% confidence interval of the regression was then extracted for each stage. Linear regressions from different intervals were compared via ANCOVA (an F-test) (Keppel, 1991; Lomax, 2007; Montgomery, 2012; Tabachnick and Fidell, 2007). A quadratic fit was also tested and Bayesian Information Criteria (BIC) (Dziak et al.; Wit et al., 2012) was calculated to compare fits for preference over both the core temperature range and entire viable temperature range.
Shortly identical analysis was conducted on frog videos as on fly videos above. Differing from above stage times were calculated in relation to 3$^{rd}$ cleavage (time zero) and over the assumed core linear range of 12.2 to 25.7 $^{o}$C.

#### Simulations of sequential multi-reaction networks

A Global Optimization was performed, using the fmincon function and MultiStart in matlab, to maximize the curvature of our sequential linear reaction series equation (E1). Optimization was done for 2 reactions with constraints on $E_a$ (20-100 kJ (Lepock, 2005)) and on k (1 sec to 3 days). The resulting $E_a$ and A for the 2-reaction combination with highest curvature was expanded to a 1000 reaction equivalent by expanding the lower $E_a$ reaction to a 999 reaction equivalent with adjusted prefactors.

#### GAPDH activity assay

We measured GAPDH activity at various temperatures similarly to as previously described (Krebs, 1955; Velick, 1955). GAPDH from rabbit muscle was purchased from Sigma (G2267). The assay buffer contained 150 mM sodium phosphate (adjusted to pH 8.5 with HCl), 300 mM sodium arsenate, 7.5 mM NAD, 15mM DL-glyceraldehyde-3-phosphate, 3 mM DTT, 0.3 ug/mL of GAPDH. Absorbtion of 2.95 mL of this solution was preincubated for 5 minutes at the given temperature (controlled with a peltier thermostatted cell holder) and absorption measured at 340nM in a quartz cuvette in an Agilent Cary 300 spectrophotometer. Samples were continuously mixed with magnetic stir bar. After pre-incubation, we added 50uL of 15mM D/L glyceraldehyde-3- phosphate (Caymen) and continuously monitored absorption at 340 nM for at least five minutes.
